# Navigating the Fitness Landscapes of Plasmodium falciparum Dihydrofolate Reductase: Evolutionary Insights into Antifolate Resistance

**DOI:** 10.1101/2024.10.31.621387

**Authors:** Danny Muzata, Dwipanjan Sanyal, A Shivram, Deeptanshu Pandey, Soumyananda Chakraborti, Aneesh S. Chivukula, Ravi Jasuja, Vladimir N. Uversky, Daniel Zielinski, Prajna D. Upadhyay, Mark Albers, Sourav Chowdhury

**Affiliations:** One Health Trust, Bangalore, India; Department of Computer Science and Information Systems, Birla Institute of Technology and Science-Pilani, Hyderabad, India; Structural Biology and Bioinformatics Division, CSIR-Indian Institute of Chemical Biology, Kolkata, India; Department of Biological Sciences, Birla Institute of Technology and Science-Pilani, Hyderabad, India; Research Program in Men’s Health: Aging and Metabolism, Brigham and Women’s Hospital, Harvard Medical School, Boston, Massachusetts, USA; USF Health Byrd Alzheimer’s Research Institute, Morsani College of Medicine, University of South Florida, Tampa, FL 33612, USA; Department of Bioengineering, University of California, San Diego, La Jolla, CA, USA; Department of Neurology, Massachusetts General Hospital, Harvard Medical School, Boston, Massachusetts, USA

**Author notes:** Corresponding Author: Sourav Chowdhury, Department of Biological Sciences, Office: J Block-109, Birla Institute of Technology and Science, Hyderabad Campus, Hyderabad, Telangana, India, 500078. These authors contributed equally to the work (Danny Muzata, Dwipanjan Sanyal).

**Keywords:** Malaria, Plasmodium falciparum, Sequence space analysis, Protein fitness, Dihydrofolate reductase, Protein evolution, Antifolate resistance, Folate pathway, Drug design

## Abstract

The rapid emergence of drug resistance in malaria parasites poses a significant challenge to the efficacy of antifolate treatments. Traditional drug development approaches, which often rely on empirical screening with limited mechanistic insights, tend to overlook the complex evolutionary mechanisms that enable *Plasmodium falciparum* to evade drug inhibition while preserving enzyme functionality. In this study, we employed computational techniques to investigate the mutational landscape of dihydrofolate reductase (DHFR), focusing on regions essential for enzyme stability and resistance. Our analysis uncovered conserved residues essential for stability, mutation hotspots that enhance adaptability under drug pressure and co-evolving clusters revealing critical functional interdependencies. Through integrated approaches including mutational scanning, epistatic interaction modeling, and fitness trajectory mapping, we elucidated distinct evolutionary pathways that drive resistance. We were able to track the adaptive paths taken by wild-type residues upon mutation, revealing the steps required to reach high-fitness peaks within the rugged fitness landscape. These findings provide valuable insights into the molecular mechanisms of antifolate resistance. We suggest that future drug design should target co-evolving networks and conserved regions to support the development of next-generation therapies to overcome resistance.

## Introduction

To date, malaria remains one of the most challenging infectious diseases to control partly due to the parasite’s ability to rapidly evolve resistance to frontline drugs^1,2^. Among the most challenging cases of resistance are mutations in *P. falciparum* dihydrofolate reductase (*pf*DHFR), which reduce the effectiveness of antifolate therapies^3,4^. Antifolate drugs have long been used in treating not only malaria but other diseases including bacterial infections and cancer^5,6^. These drugs exert their effects by targeting and inhibiting key enzymes in folate metabolism, a critical pathway for the synthesis of nucleotides and certain amino acids. In *P. falciparum*, several metabolic pathways are targeted by antimalarial drugs. One good example is the hemoglobin degradation pathway and the mitochondrial electron transport chain that is targeted by drugs such as chloroquine and artemisinin^2^. Heat shock proteins also present promising drug targets due to their essential roles in parasite survival and virulence across the various life cycle stages^1^. These alternative drug targets highlight the complexity of *P. falciparum* biology and emphasize the importance of targeting multiple pathways to combat resistance. However, the folate pathway, particularly DHFR, remains one of the most critical targets in antimalarial therapy due to its central role in DNA synthesis and repair^7–9^.

In *P. falciparum*, DHFR catalyzes the NADPH-dependent reduction of dihydrofolate to tetrahydrofolate, an essential cofactor required for the synthesis of purines, thymidylate, and certain amino acids^8,10^. This reduction reaction makes DHFR an indispensable enzyme in the folate pathway and a prime target for antifolate drugs such as pyrimethamine and cycloguanil^8,11^. Single and multiple mutations at residue positions 16, 50, 51, 59, 108, and 164 of *pf*DHFR have been linked to antifolate resistance in malaria. Mutations at these sites confer resistance to pyrimethamine and cycloguanil by reducing its binding affinity to the protein rendering the drug less effective^3,12,13^.

Despite significant research into DHFR mutations and their role in antifolate resistance, gaps remain in understanding how these mutations affect protein fitness, structure, and function at the molecular level. The concept of sequence space provides a theoretical framework for mapping all possible amino acid combinations in a protein and exploring the evolutionary constraints that shape functional regions. Sequence space analysis has proven valuable in uncovering evolutionarily constrained regions as well as identifying conserved and flexible sites within protein structures^14^. Thus, in this study, we carried out a thorough sequence space analysis of DHFR protein to explore the evolution of antifolate resistance and its implications for drug design. We developed an evolutionary dynamics simulation model to map adaptive fitness trajectories. This approach integrates both deterministic and stochastic elements, simulating how mutations traverse the fitness landscape while balancing short-term gains and long-term evolutionary potential.

## Results

### Sequence space analysis

In this study, we explored the sequence space of *pf*DHFR to gain insights into how mutations influence protein fitness and stability. This sequence-based study provided important insights into the protein’s evolutionary dynamics. It highlighted how mutations at critical positions influence its stability, function, and mutational tolerance. Through careful analysis of sequence conservation, co-evolution and network-based interactions, we inferred how sequence variability could lead to significant functional consequences in the enzyme, especially in the context of resistance-related mutations.

### Shannon entropy analysis revealed critical positions essential for protein function and mutation-tolerant positions

We performed Shannon Entropy (SE) calculation for each residue position to identify conserved and variable regions within the protein sequence which play a crucial role in the proteins structure and function. Our analysis revealed six conserved residue positions, that is, positions 30, 32, 38, 39, 91 and 104 which exhibited zero entropy values (**Figure 1A**). These regions are likely to be evolutionarily constrained, as mutations in these positions could disrupt protein stability or function. Certain active site positions were also found to be relatively conserved (close to zero) implying that mutations in these positions could have disruptive effects on the pyrimethamine drug binding. Conversely, regions with higher entropy values were identified mostly in the flexible loop regions between residue positions 9–25 and 69–89. Other regions towards the C-terminus (190–219) also displayed high spikes (**Figure 1A**). These positions displayed greater variability, suggesting they may be more tolerant to mutations. Positions C41, N42, C50, and S99 also exhibited high variability and have mutations linked to cycloguanil and pyrimethamine resistance.

**Figure 1:**
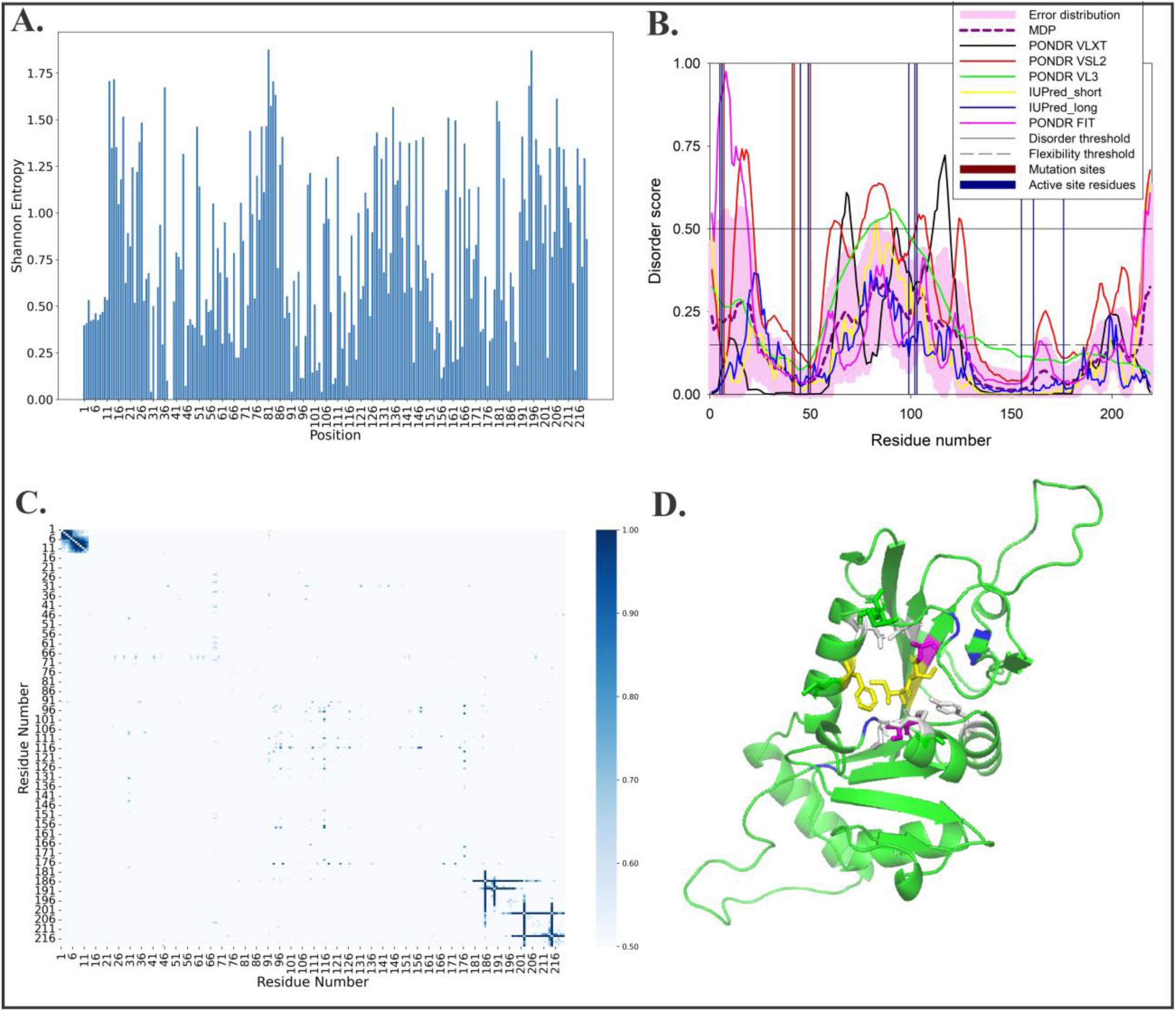
Sequence Space Analysis of DHFR Reveals Key Regions Under Evolutionary Constraints and Functional Adaptation. **(A)** Shannon Entropy plot shows sequence variability across the DHFR domain. Residues with higher Shannon Entropy scores exhibit greater sequence variability (high spikes), while residues with lower scores (close to zero) are highly conserved. The mutation sites *C41*, *N42*, *C50*, and *S99* exhibit high entropy, indicating their propensity for evolutionary changes. **(B)** Per-residue disorder propensity of pfDHFR evaluated by six commonly used disorder predictors: PONDR® FIT, PONDR® VSL2, PONDR® VL3, PONDR® VLXT, IUPred Short, and IUPred Long. MDP corresponds to the mean disorder profile calculated by averaging the outputs of individual predictors. Residues with disorder scores ≥ 0.5 are considered disordered, while those between 0.15 and 0.5 are flexible. **(C)** The Mutual Information (MI) plot illustrates strong MI values (darker blue) indicating residues that co-vary. **(D)** 3D structure of the DHFR protein, color-coded by regions of variability and highlighting mutation and active sites. Active site residues are highlighted in yellow sticks, while mutation sites are highlighted in magenta sticks. High-entropy regions in active site are shown in white indicating areas prone to evolutionary change. Highly conserved regions are shown in blue.

### Intrinsically disordered prediction revealed that the DHFR is predominantly ordered

We further explored the structural flexibility of the DHFR using six commonly used disorder predictors^15^: PONDR® FIT^16^, PONDR® VSL2^17^, PONDR® VL3^17^, PONDR® VLXT^18^, IUPred Short^19^, and IUPred Long^19^ and generated its disorder profile. The outputs of these predictors were averaged into a mean disorder profile (MDP), providing a consensus view of the protein’s disorder propensity. Intrinsic disorder often correlates with regions capable of tolerating mutations contributing to evolutionary flexibility.

Our findings revealed that the DHFR protein is predominantly ordered, with most residues displaying low disorder scores (< 0.5), indicating structural stability. A few flexible regions (disorder scores between 0.15 and 0.5) were identified at positions 1–24, 60–125, 199–203, and 214–219. However, as per the outputs of PONDR® VSL2^17^, which is one of the more accurate disorder predictors, several key regions of elevated disorder were identified, that is, residues spanning position 13-21, 61-65, 76-90, 103-107, 123-124, and 217-219. These regions suggest points of potential mutational tolerance, contributing to the protein’s adaptability (**Figure 1B**).

### Mutual information calculation revealed tight clusters of inter-dependent residues in the N-terminus and C-terminus

While SE calculation offered insights into evolutionary conservation rates, it did not account for interactions between individual residues, particularly how certain positions may evolve in tandem due to structural or functional dependencies. To explore these interactions, we employed Mutual Information (MI) analysis. This measures the co-evolution of residue pairs and identifies compensatory mutations that help maintain protein structure and function. The MI analysis revealed key co-evolutionary relationships between specific residues, providing critical insights into the structural and functional constraints imposed by selective pressures. A high MI value indicates significant positional interdependence, reflecting strong co-evolutionary relationships between residues. In the MI heatmap (**Figure 1C**), dense clusters, such as those seen between residues 1–11 and 181–216, highlight regions with notable co-variation.

### MI interaction networks highlight co-evolution of mutational sites in DHFR sequence space

Building on the MI calculations, the graph-based analysis was performed to vividly reveal the complex co-evolutionary relationships between residues in the active site. These MI networks provide a clear visual representation of how different regions of the protein interact. The residue-residue networks provided detailed evidence of epistatic interactions, where mutations at one site could influence compensatory changes at another, preserving enzyme function. The interaction networks for I5 and C6 (**Figure 2A**) demonstrated a tightly knit structural cluster potentially critical for drug-binding and resistance mechanisms. They exhibited strong interactions (MI > 0.8) with residues positions C8, C9, K10, V11, K19, and E12. Residues F49 and I103 also displayed strong and moderate interactions with M95, I103 and K124, S200, F107, and L110 respectively, suggesting their central role in maintaining the active site configuration.

**Figure 2:**
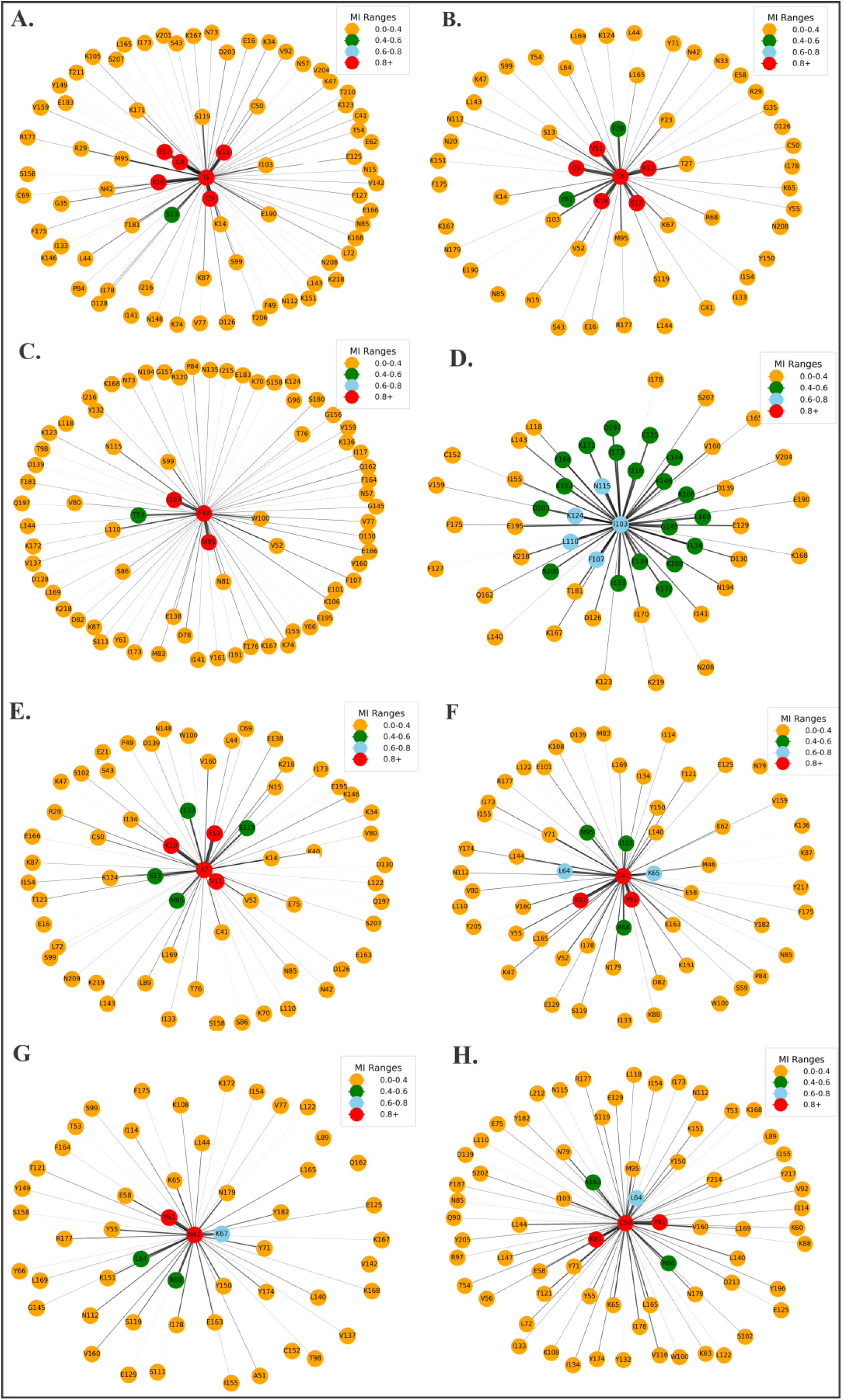
Mutual Information Networks Reveal Co-evolving Residues Critical for DHFR Function and Adaptation. (**A**) Interaction network for active site **I5**, showing strong MI values with residues like I2, C8, and D1. (**B**) This shows a network centered on active site **C6** with strong interactions extending to K10, V11, and C9, implying that these residues may co-evolve to maintain catalytic function under selective pressures. (**C**) Interaction network for **F49** showing significant interactions with residues such as I103 and T53. (**D**) Interaction network around active site **I103** displayed extensive interactions with several residues such as N115 and K124 highlighting the interconnectedness of residues involved in DHFR’s active site dynamics. (**E**) Interaction network centered on mutation site **A7** showing significant correlations with residues such as V11, A4, and K10. (**F**) Network around mutation site C41, displaying strong MI interactions with residues like K55 and C9. (**G**) Mutation site N42 shows strong interactions with W103 and K67 (**H**) Network focused on mutation site C50 highlights strong interactions with residues K55 and V11.

We extended this analysis to the mutational sites which also revealed intriguing co-evolutionary relationships. Specifically, residues like A7 showed significant interactions with K10, V11 and E12 while C41 interacted strongly with Y61 and K67. Similarly, N42 and C50 formed interactions with Y61 and K67.

### Maximal clique analysis and community detection identified densely connected cliques

In our study, we performed Maximal Clique Analysis (MCA)^20,21^ and Community Detection^22^ to identify densely connected sub-networks (cliques) and partition the protein into distinct interaction groups. These analyses provide a deeper layer of insight compared to MI analysis, as they go beyond pairwise interactions by revealing how clusters of residues interact within a broader network. Important cliques were identified in regions containing known active and mutation sites, including residues 6, 7, 42, 49, 155 (**Figure 3A**). These residues form part of highly interconnected clusters, indicating strong structural and functional interdependencies.

**Figure 3:**
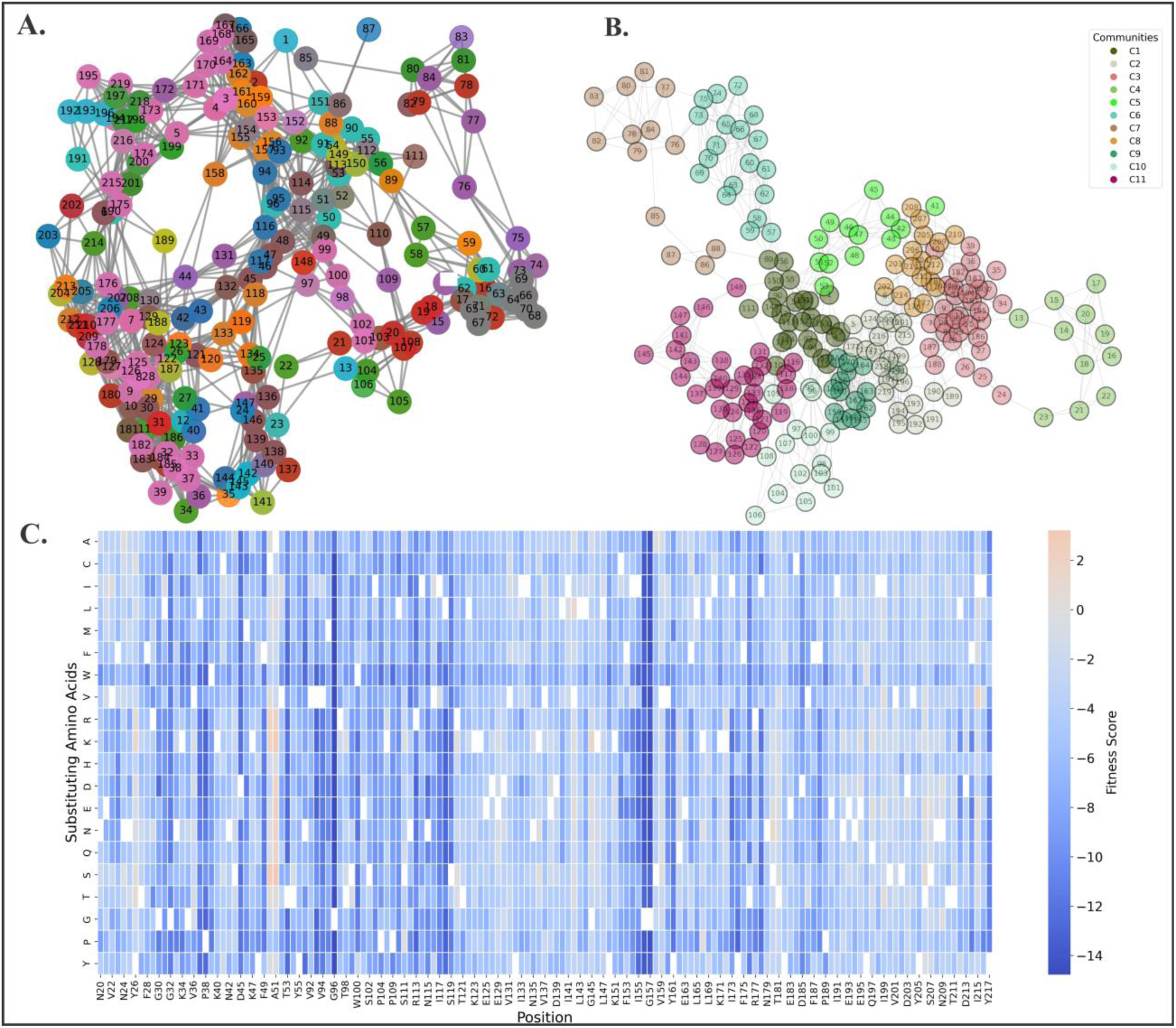
Maximal Clique and Mutational Sensitivity Analyses Reveal Inter-Residue Networks and Evolutionarily Constrained Regions in DHFR. (**A**) Maximal clique detection analysis visualizes cliques or complete subgraphs within the DHFR residue network. Different cliques represent tightly connected regions of residues which indicate areas of high inter-residue coordination and co-evolution. (**B**) Community detection analysis using a modularity-based algorithm identified 11 distinct communities within the DHFR protein. These communities represent clusters of residues that are likely to have high intra-group connectivity. (**C**) Deep mutational scan heatmap showing the fitness effects of single amino acid substitutions across the DHFR protein. The heatmap reveals positions that are highly sensitive to mutations, with dark blue indicating deleterious effects (negative fitness scores) and light shades or orange representing beneficial or neutral mutations.

Residues 49 and 155, which form part of pyrimethamine-binding pocket, were observed in multiple cliques alongside other critical residues highlighting their importance in stabilizing the surrounding structural framework of the protein. Mutation sites such as residues 7 and 42 also formed part of distinct cliques where they interacted with other residues. It is worth noting that residue 6, a drug-binding pocket, was identified within the same clique as residue 7, a known mutation site. The analysis revealed that the active sites and mutation sites often reside in highly interconnected cliques, suggesting that mutations at these sites might affect the stability and function of nearby residues through their cooperative interactions. Community detection (**Figure 3B**) further uncovered broader groupings of residues that frequently interact within the same sub-network. It highlighted functional domains or regions that evolve together.

### Deep mutational scanning yielded mutational hotspots with beneficial fitness effects that may facilitate evolutionary adaptation under drug pressure

The Deep Mutational Scanning (DMS) analysis was performed to gain insights into the functional consequences of amino acid substitutions across the protein sequence. By quantifying the fitness effects of each mutation, this analysis allowed us to map the sequence space of DHFR, identifying regions that are intolerant to mutations, as well as those that can tolerate or even benefit from substitutions. A significant number of deleterious mutations were observed in key regions of the DHFR enzyme (**Figure 3C**), particularly around the active site. Positions such as I155, which is a critical residue within the active site, show dramatic reductions in fitness for nearly all substitutions. This suggests that I155 plays an essential role in maintaining the enzyme’s catalytic function and structural stability. Similarly, D45, F49 and I155, another set of residues located in the active site, also shows a strong deleterious signal upon mutation with almost all substitutions. The positions G96, G156, G157 and Y217 strongly emerged as key sites where mutations lead to significant fitness losses. These residues are situated near the catalytic core of the enzyme and substitutions here likely disrupt the enzyme’s stability or interfere with essential catalytic processes. The DMS analysis also uncovered key mutation hotspots with beneficial fitness effects, particularly in the regions spanning residues 120 to 155 and 190 to 211.

### The fitness trajectories for active site residues illustrates distinct adaptive paths for various mutations, highlighting the complexity of the fitness landscape for pfDHFR

Our fitness trajectory analysis provides critical insights into the distinct adaptive pathways of key active site residues in *pf*DHFR. This reveals how the enzyme navigates complex fitness landscapes under selective pressure. To simulate these evolutionary paths, we developed an epistasis-based fitness prediction model using Dijkstra’s shortest path algorithm. In this model, mutations are represented as nodes, and the edges between them are weighted by the absolute fitness differences between the mutations. This approach identifies the most efficient mutational paths—those requiring the fewest steps—from the wild-type to both local and global fitness peaks. The model captures both stochastic and deterministic evolutionary dynamics, offering a realistic framework for simulating biological evolution. Stochastic sampling introduces randomness, reflecting genetic drift and the exploration of multiple paths. Deterministically, the algorithm favors mutations with immediate fitness benefits, emulating natural selection. This dual approach balances short-term incremental adaptation with long-distance evolutionary leaps, mirroring the unpredictable yet selective nature of real-world evolution.

Using deep mutational scan data to guide these pathways, the model reveals how near-neutral and compensatory mutations expand the genotype-phenotype space. This allows the enzyme to smooth rugged fitness landscapes and adapt under selective pressure. For instance, mutations at position 49 exhibited quick transitions to fitness peaks such as A51S, highlighting the mutational tolerance at this site. However, other substitutions at the same site, such as F49K and F49W, required longer pathways to reach comparable peaks, indicating variability in their fitness effects (**Figure 4A**). Similarly, mutations S99C and S99T demonstrated shorter evolutionary paths to local fitness peaks like C50S, suggesting faster adaptation, while mutations like S99F followed more extended trajectories (**Figure 4C**). This trend was also observed at positions 102 and 103, with certain mutations achieving rapid adaptation, while others required more gradual, complex walks. Conversely, position 155 presented one of the most challenging landscapes, with most mutations demanding over 100 evolutionary steps to achieve fitness peaks (**Figure 4F**). A comparable trend was seen at position 161 (**Figure 4H**), where mutations such as Y161L and Y161K required long evolutionary walks toward global peaks, highlighting the ruggedness of the landscape at these critical sites.

**Figure 4:**
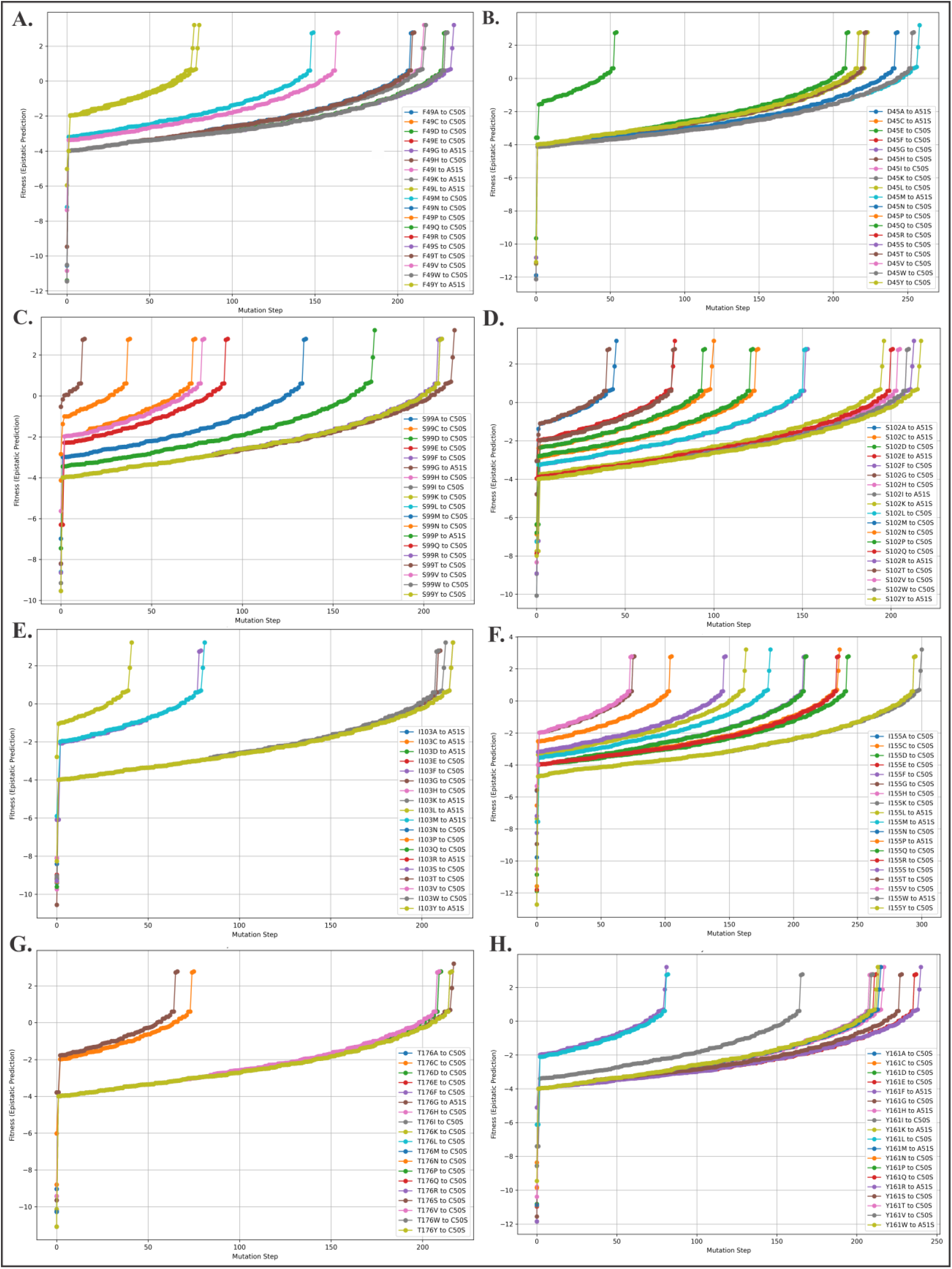
Fitness Trajectories Highlight Adaptive Paths of Key Active Site Residues in DHFR. **(A)** At position 49, mutations such as F49L and F49Y achieve quicker routes to fitness peaks, while others like F49K, F49W, and F49H follow longer, more gradual trajectories. **(B)** Position 45 shows gradual fitness increases across all mutations, except D45E which achieved a relatively quicker fitness. **(C)** Mutations at position 99 exhibit variability in trajectory length. Positions S99T and S99C reach fitness peaks faster, while S99G requires the longest path. **(D)** Position 102 shows variation in mutation paths, with S102A and S102P achieving fitness improvements more quickly than S102E and S102F. **(E)** Mutations at positions 103 generally follow longer trajectories to fitness peaks. **(F)** At position 155, most mutations require extended walks of over 100 steps to reach fitness peaks. **(G)** Shows similar patterns at position 161, where Y161L and Y161K mutations follow longer trajectories. **(H)** Position 176 demonstrates both short and long trajectories, with some mutations reaching fitness peaks more rapidly, while others require extended adaptive steps.

These findings underscore that evolutionary adaptation in DHFR is not uniform. While certain residues, such as 49 and 99, serve as stepping stones toward resistance by facilitating quick fitness gains, others like 155 demand more extensive evolutionary navigation. The interplay between stochastic exploration and deterministic selection captured by our model reflects the intricate dynamics of DHFR evolution, offering a deeper understanding of how the enzyme balances adaptability and functional constraints under selective pressure.

## Discussion

The folate pathway plays a crucial role in the survival of *P. falciparum* by supplying essential cofactors for nucleotide and amino acid synthesis^7^. This makes DHFR a prime target for antifolate drugs. However, the emergence of resistance mutations in this essential protein has led to significant challenges in the effective treatment of malaria. These mutations have enabled the parasite to evade drug inhibition while preserving enzyme functionality^23^. Further complexity arises from the genomic organization of the *pfdhfr-ts* gene. This is a bifunctional gene located on chromosome 4 that encodes both DHFR and thymidylate synthase within the same polypeptide chain^24,25^. This dual functionality introduces potential pleiotropic effects, where mutations selected for resistance in the DHFR domain could unintentionally affect the function of the TS domain, or vice versa. Similar to the observations of Raff and Kaufman^26^, such interdependencies highlight the complexity of interconnected biological systems, where targeted changes can yield large-scale, unintended consequences across multiple functions. The interdependence between these two enzymatic activities creates challenges in predicting the evolutionary consequences of mutations, as adaptations beneficial for one activity might impair the other. While the direct impact of promoter mutations on this gene remains largely unexplored^27^, these mutations could potentially influence gene dosage and expression levels, thereby affecting drug resistance mechanisms. This genomic context implies that resistance arises not only from individual enzyme mutations but also from the complex regulation of gene expression. Thus, understanding the evolutionary dynamics driving these mutational escapes in DHFR is essential for designing novel antifolate therapies.

Mutational landscapes are inherently complex^28^, reflecting the diverse roles amino acid substitutions play in shaping protein function and evolution. Proteins, including DHFR, are subjected to a range of evolutionary pressures that necessitate both structural stability and adaptability^29,30^. However, mutations that confer drug resistance often introduce trade-offs that impact their fitness^31^. This balance is complicated by the fact that near-neutral mutations, while individually minor, can act as stepping stones toward significant adaptive shifts^32^. The study of mutational landscapes using tools such as deep mutational scanning has proven critical in identifying mutations that influence protein adaptability without drastic functional compromises^33,34^. Some mutations improve fitness only marginally but, over time, accumulate to confer drug resistance^35^. These insights highlight how proteins like DHFR navigate a rugged landscape, with regions of both mutational tolerance and structural constraint. This understanding is essential for predicting the evolution of resistance and identifying critical points for therapeutic intervention.

The evolutionary landscape of proteins is defined by peaks of high fitness separated by valleys of low fitness. This ruggedness reflects the complex interplay between mutations, structural integrity, and functional performance. Adaptive walks along this landscape often involve both stochastic exploration and deterministic processes that select for advantageous traits^36^. Near-neutral mutations facilitate exploration by expanding the search space which allows enzymes to navigate fitness valleys more efficiently^32^. Over time, these mutations can accumulate to provide a selective advantage, especially under environmental pressures. Evolutionary trajectories are further shaped by epistasis, where interactions between multiple mutations influence fitness outcomes^37^. These complexities emphasize the importance of studying not just individual mutations but also their collective impact on protein function.

The present study provides critical insights into the mutational and evolutionary landscapes of *P. falciparum* DHFR with emphasis on both structural constraints and adaptive strategies employed by the protein. Conserved residues were identified and highlighted as structurally indispensable elements that are less tolerant to mutations. These regions offer promising targets for novel antifolate therapies, as disrupting them could impair the parasite’s ability to maintain functionality and evolve resistance. Network-based analyses further revealed key hubs and cliques of interacting residues. The presence of such interconnected, co-evolving clusters highlights that resistance arises not through isolated mutations but via cooperative networks that preserve stability despite structural alterations. Several mutation hotspots were also identified with beneficial fitness effects, particularly in regions spanning residues 120–155 and 190–211. While many mutations, especially at key active site residues like I155, D45, and F49, caused significant fitness losses due to their critical role in protein stability, certain regions exhibited greater tolerance to substitutions. These mutation-tolerant regions suggest potential adaptive flexibility which enable the enzyme to navigate selective pressures by accommodating beneficial changes without compromising overall functionality. The integration of conserved elements, mutation hotspots, co-evolving clusters, and interconnected networks underscores the multifaceted nature of resistance. Targeting these networks and conserved regions provides a promising approach for the development of future-proof antifolate therapies, disrupting the parasite’s ability to adapt and evolve in response to drug pressure.

To simulate adaptive evolutionary pathways and explore how mutations influence fitness landscapes, we developed an epistatic fitness prediction model. Our model provides valuable insights into how *P. falciparum* DHFR navigates complex fitness landscapes under selective pressure. This dual framework captures the unpredictable aspects of evolution while also reflecting selective pressures that favor beneficial mutations. Through this approach, we modeled both rapid adaptive walks and gradual evolutionary trajectories, mirroring the diverse paths proteins take toward higher fitness in dynamic environments. The use of deep mutational scan data enabled us to assign fitness values to individual mutations. This provided a quantitative lens to evaluate how various mutations affect enzyme stability and function. By mapping these trajectories, we demonstrated that near-neutral mutations and compensatory interactions play pivotal roles in expanding the genotype-phenotype space, thereby smoothing the rugged fitness landscape. This adaptive flexibility allows DHFR to navigate resistance pathways efficiently, overcoming mutational barriers while retaining functionality.

The insights from this model emphasize the importance of epistatic interactions—where mutations at one site influence the fitness outcomes of others—in shaping evolutionary pathways. It underscores that resistance in DHFR is not merely a product of isolated mutations but rather emerges from complex mutational networks that facilitate both short-term gains and long-term adaptability. This predictive framework enhances our understanding of antifolate resistance mechanisms and lays the groundwork for designing novel antifolate therapies that can effectively disrupt these evolutionary strategies.

While our study offers significant insights into the evolution of antifolate resistance, it is not without limitations. This research was solely based on protein sequence analysis and did not delve into the structural dynamics of DHFR, which could provide additional insights into enzyme stability, conformational changes, and protein-drug interactions. Despite incorporating stochastic sampling to simulate randomness in evolutionary choices, the model may still miss certain complexities of real-world adaptation, such as temporary fitness losses or parallel exploration of multiple paths. These limitations highlight the challenges of representing rugged fitness landscapes, where local optima may be revisited and evolutionary trajectories can follow non-linear patterns over time. Moreover, the limited availability of fitness data constrains the scope of the fitness landscape explored. Expanding the dataset or integrating complementary experimental data would provide a more comprehensive view of mutational effects across a broader range of residues. Future experimental validation of our computational findings is crucial. Molecular docking simulations and fitness assays in biological systems will confirm whether the identified adaptive pathways correspond to real-world resistance mechanisms. Integrating advanced computational tools, such as machine learning models, could enhance predictive accuracy and uncover novel mutational pathways that are difficult to identify using conventional algorithms. These efforts will contribute to the development of future-proof antifolate therapies capable of addressing the challenges posed by evolving resistance in *P. falciparum*.

## Materials and Methods

### Sequences preprocessing

For this study, we used the crystal structure of the wild-type *P. falciparum* DHFR-TS protein (PDB ID: 3QGT) as the reference for our analyses. The protein structure consists of two domains: the DHFR domain (first 231 residues) and the TS domain (last 288 residues), separated by an 89-residue interjunction region (Supplemental Figure 1). Given the focus on DHFR, we truncated the DHFR domain, which stretches from amino acid position 10 to 228 in the full-length protein as annotated in UNIPROT^38^. This truncation was performed using the PYMOL Molecular Graphics System (version 3.0.3)^39^. For consistency, the truncated sequence was re-indexed from 1 to 219, a scheme used throughout the study.

### Homology Modeling and Structure Validation

Upon examination, the DHFR protein contained ten missing residues in its sequence from position 86 to 95. To fill in these gaps, homology modeling was performed using Modeller (Version 10.5)^40,41^. This software generated several structural models and ranked them using the Discrete Optimized Protein Energy (DOPE) scoring function. The model with the lowest DOPE score was selected as the most accurate representation of the full DHFR structure (**Supplemental Figure 2**). To ensure the accuracy of the modeled structure, we calculated the Root Mean Square Deviation (RMSD) between the modeled residues and the crystal structure (**Supplemental Figure 3**). An RMSD value of 0.306 Å indicated high structural similarity. After validation, we extracted the amino acid sequence (219 residues) from the modeled DHFR structure for further analyses.

### Sequence Space Analysis

To explore the mutational landscape of DHFR, we performed a sequence space analysis using the multiple sequence alignment (MSA) of homologous DHFR proteins. First, we retrieved 5000 non-redundant protein sequences homologous to DHFR using the NCBI BlastP tool^42^. The sequences were cleaned to ensure high quality, non-redundancy and biological relevance. We then aligned the cleaned sequences using the ClustalW algorithm^43^, and the resulting MSA served as the basis for the subsequent analyses.

### Occupancy Calculation

Occupancy calculations were performed to assess how frequently specific amino acids occupy each residue position in homologous DHFR sequences. Positions with high occupancy values represent conserved residues, likely essential for the protein’s structural stability. By comparing occupancy data with mutation sites, we identified residues where mutations may have significant impacts on the protein’s structure and function.

### Shannon Entropy Calculation

Shannon entropy (S(*i*))^44^ was calculated to quantify the variability of amino acids at each sequence position. Positions with low entropy values represent conserved residues, critical for structural and functional integrity, while high entropy values indicate positions prone to mutations.

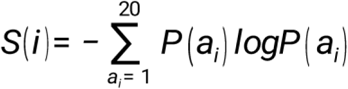

Where ***i*** represents the sequence position, and ***P(a_i)*** is the probability of amino acid ***a*** occurring at position ***i*** in the MSA. Lower values of ***S(i)*** correspond to highly conserved residues, while higher values suggest increased variability.

### Mutual Information Calculation

To investigate co-evolving residues, we calculated mutual information (MI)^45,46^ for each pair of positions in the MSA. MI quantifies the degree of correlation between two sequence positions, revealing residues that might be functionally or structurally interdependent. These co-varying residues could indicate regions where mutational compensation occurs, maintaining overall protein function despite mutations in specific positions.

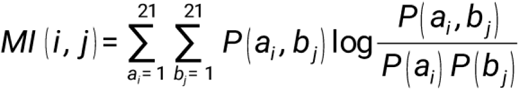

In this formula, *P(ai, bj)* represents the probability of amino acids a and b occurring at positions *i* and *j*, respectively, while *P(a_i_)* and *P(b_j_)* are the individual probabilities at each position. The *MI(i,j)* values range from 0 (uncorrelated residues) to *MI_max_* (the most interdependent residue pairs). In this study, a cutoff MI value of 0.5 was chosen to focus on highly co-varying positions. The MI values were visualized as a heatmap and a co-evolutionary matrix, generated using python programming.

### Intrinsic Disorder Prediction

Intrinsic disorder prediction analysis was conducted to explore the presence of disordered regions within the DHFR protein. Disordered regions, known for their inherent flexibility, can tolerate higher levels of mutation and often play significant roles in protein-protein interactions. For a comprehensive evaluation, we employed six widely used disorder predictors: PONDR® FIT^16^, PONDR® VSL2^17^, PONDR® VL3^17^, PONDR® VLXT^18^, IUPred Short^19^, and IUPred Long^19^. These predictors leverage distinct predictive approaches to identify disordered regions within the protein. PONDR® VSL2 is particularly optimized for proteins containing both structured and disordered regions^47^. By integrating the outputs from all six predictors, we generated a mean disorder profile (MDP), providing a consensus disorder score for each residue. Residues with scores above 0.5 were classified as disordered, while those between 0.15 and 0.5 were considered flexible but ordered. This disorder profile was further analyzed to examine potential correlations between disordered regions, known mutation sites, and active site residues, helping to elucidate the role of structural flexibility in resistance mutations and the maintenance of protein function.

### Graph-Based Analysis

Following the MI calculations, we performed a graph-based analysis of the identified co-evolving residues. Nodes in the graph represent individual residues, while edges denote strong mutual information-based connections between them. The MI threshold was used to define the strength of these interactions, with different ranges highlighting strong (MI > 0.8), moderate (0.6 < MI < 0.8), and weak (0.4 < MI < 0.6) interactions. This graph-based analysis allowed us to visualize key mutation sites (A16, N42, C50, S99, S108, and I155) and their network of interactions, especially with active site residues. The resulting network graphs provided insight into how mutations at specific residues might influence structural stability and drug resistance through interdependencies with nearby residues.

To investigate densely connected sub-networks within the DHFR protein, we employed Maximal Clique Analysis (MCA)^20,21^ and Community Detection^22^ methods. In MCA, we aimed to identify groups of residues that form maximal cliques, where every residue (node) within a clique is directly connected to every other residue. This method focuses on identifying highly interconnected sets of residues that are structurally and functionally interdependent. Each residue in the protein is represented as a node and the edges between these nodes represent significant interactions, which are based on proximity in the protein structure or known co-evolutionary relationships. The primary goal was to identify the largest cliques within the overall interaction network. Community detection was also performed to divide the protein’s residue interaction network into distinct subgroups or communities. Unlike MCA, which focuses on fully connected sub-networks, community detection identifies groups of residues that are more likely to interact within their community than with residues outside it.

### Fitness Trajectories Analysis Using Epistatic Prediction Model

To explore the epistatic interactions between multiple mutations, we developed an epistasis-based fitness prediction model to simulate adaptive walks within the *P. falciparum* DHFR protein. The model integrates both stochastic and deterministic elements to account for the complexity of the fitness landscape. The epistatic data used were derived from deep mutation scan which predict the fitness effects of individual mutations. The goal of this analysis was to map the evolutionary trajectories of key active site residues as they navigate through the fitness landscape.

We used Dijkstra’s shortest path algorithm^48^ to construct the adaptive walks, representing the protein’s potential mutational paths in search of fitness improvement. In this graph-based approach, mutations were treated as nodes, and fitness differences between mutations as weighted edges. The shortest path corresponds to the fewest evolutionary steps required to maximize fitness. The fitness values for each mutation were calculated from the epistatic prediction data, where a positive value indicates a beneficial mutation and a negative value represents a deleterious one. Given that a set of mutations M = {m_1_, m_2_,…, m_n_} and their fitness values *f(m_i_)*, our goal was to identify the shortest adaptive walk that maximizes fitness. The distance between two mutations is defined as the absolute difference between their fitness values;

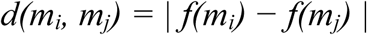

We sought to find the mutation path *P* from an initial wild-type *m_wt_* to a final high-fitness peak mutation *m_peak_*, such that the sum of fitness differences along the path is minimized;

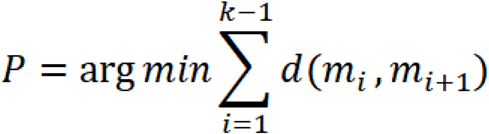

A combination of deterministic and stochastic methods was employed. The algorithm deterministically follows the mutation path with the highest fitness improvement, i.e., aiming directly for the global peak unless the local peak provides a substantially higher immediate fitness gain. When multiple peaks are available, the algorithm stochastically samples between the local and global peaks based on fitness differences. The stochastic component is controlled by a probability threshold, which adjusts based on the fitness difference between the local peak and the global peak:

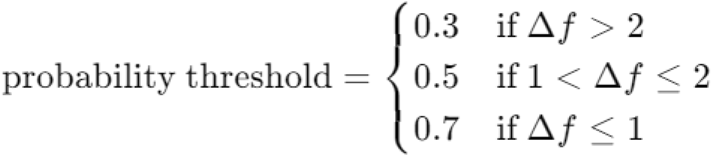

This threshold governs the likelihood of choosing a local versus a global peak, simulating a situation where evolutionary pressures might favor local exploration or long-distance leaps to distant fitness peaks.

## AUTHOR CONTRIBUTIONS

SC and DS conceived and planned the work. SC motivated and guided the work. DM performed all the formal analysis, model development, manuscript composition, and critical review. VNU performed the disorder analysis. SC, DS, VNU, SA, ASC, DZ, RJ and PDU provided critical feedback and assisted in refining the manuscript.

## ACKNOWLEDGMENT

DM acknowledges One Health Trust, Bangalore, India. DM and SC acknowledge the Director, Birla Institute of Technology and Science-Pilani, Hyderabad, India.

## DECLARATION

The authors declare that they have no conflicts of interest.

## DATA AVAILABILITY STATEMENT

The data that support the findings of this study are available from the corresponding author upon reasonable request.

## Supplemental Information

### Supplemental Figures

**Supplemental Figure S1:**
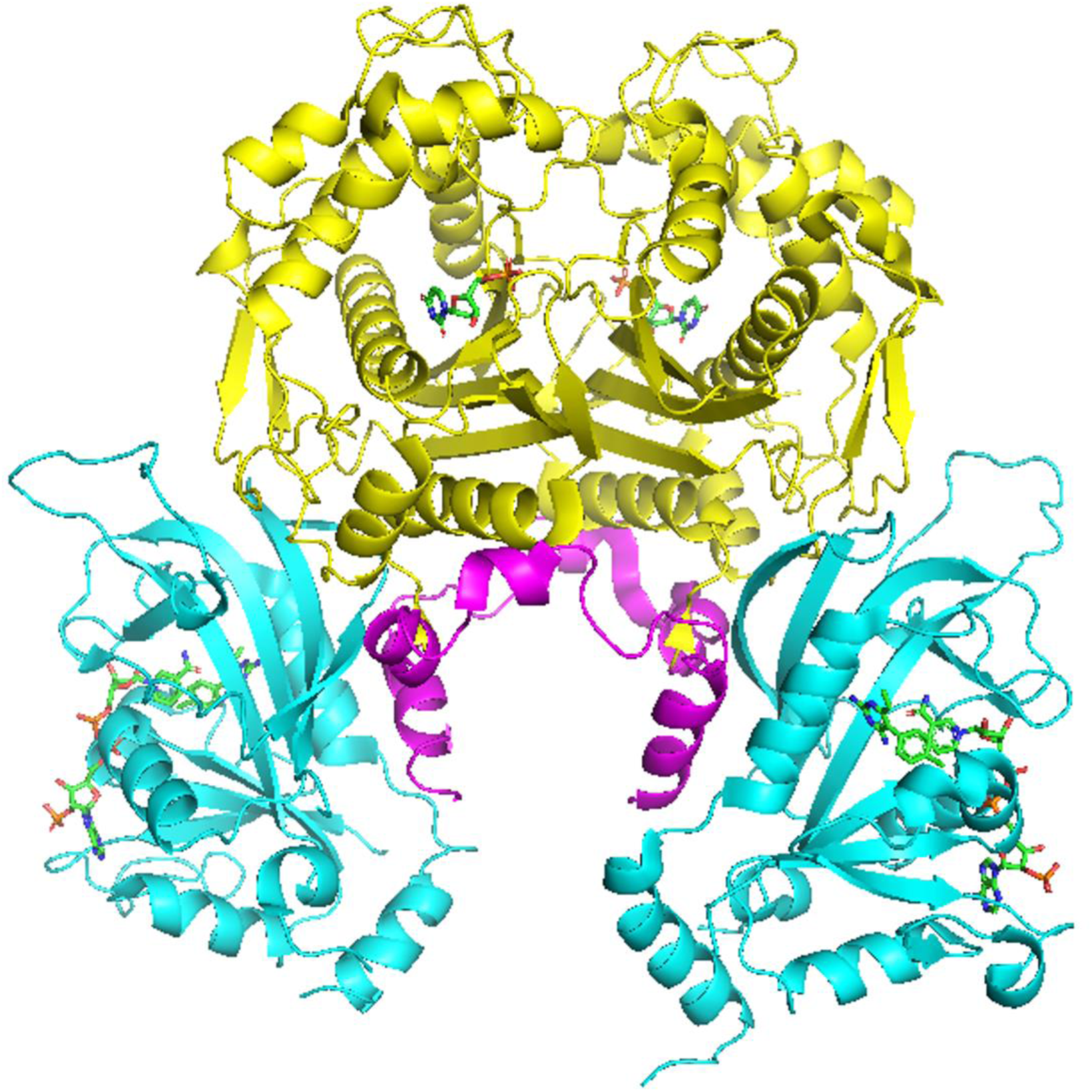
This illustrates the 608 long *pf*DHFR-TS (PDB ID: 3QGT) dimeric assembly. In cyan are the DHFR domains with DHFR-TS junction in magenta and yellow region as TS domains. The DHFR domains are complexed with the drug pyrimethamine and NADPH cofactor

**Supplemental Figure S2:**
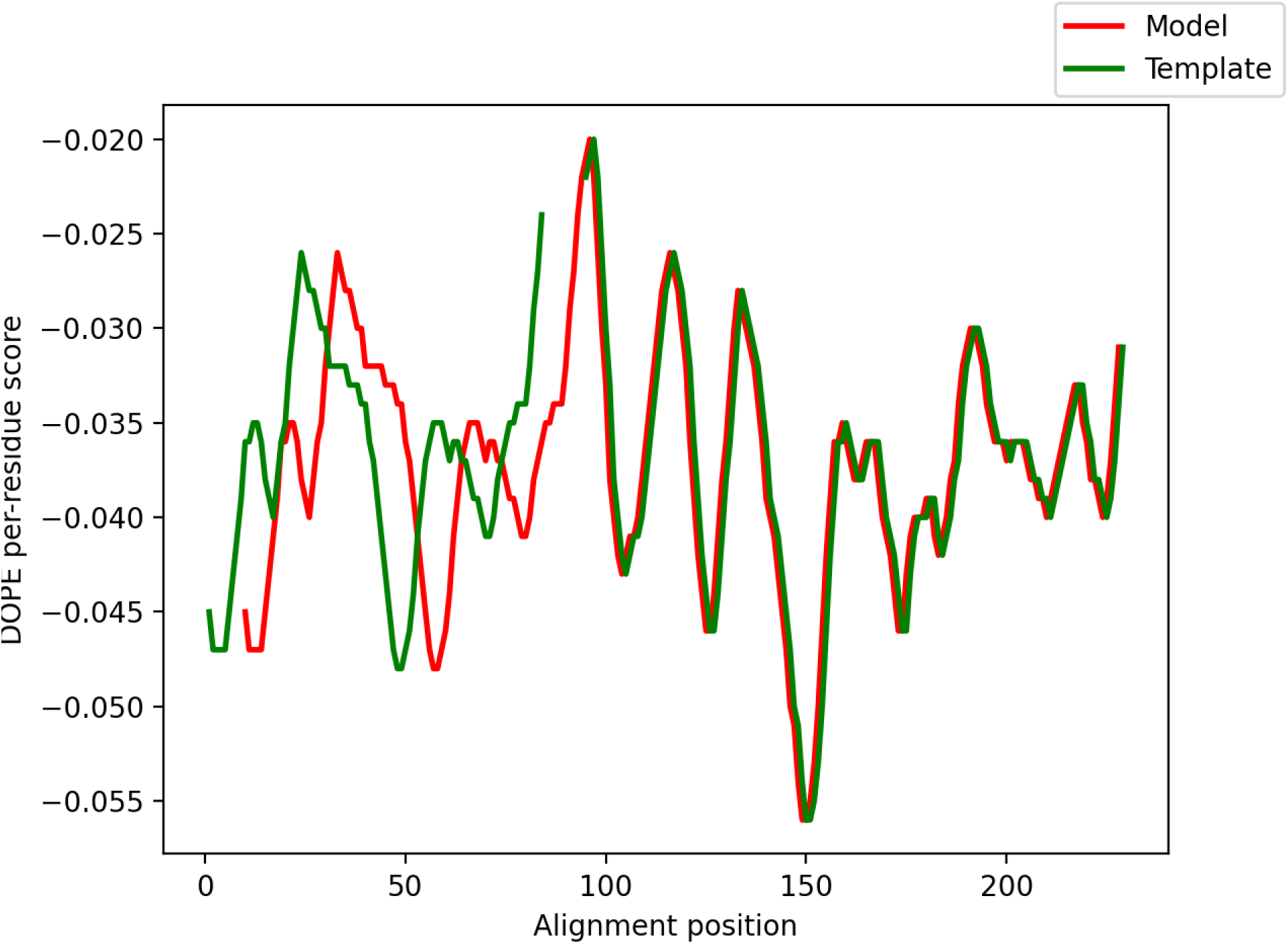
This plot presents the validation results of homology modeling performed using Modeller. The DOPE scores for the model and template structures are compared across the alignment positions. The plot shows how well the model (red) aligns with the template (green). However, the misalignment observed between positions 0 to 96 is expected as it is due to the existing gap in the template structure.

**Supplemental Figure S3:**
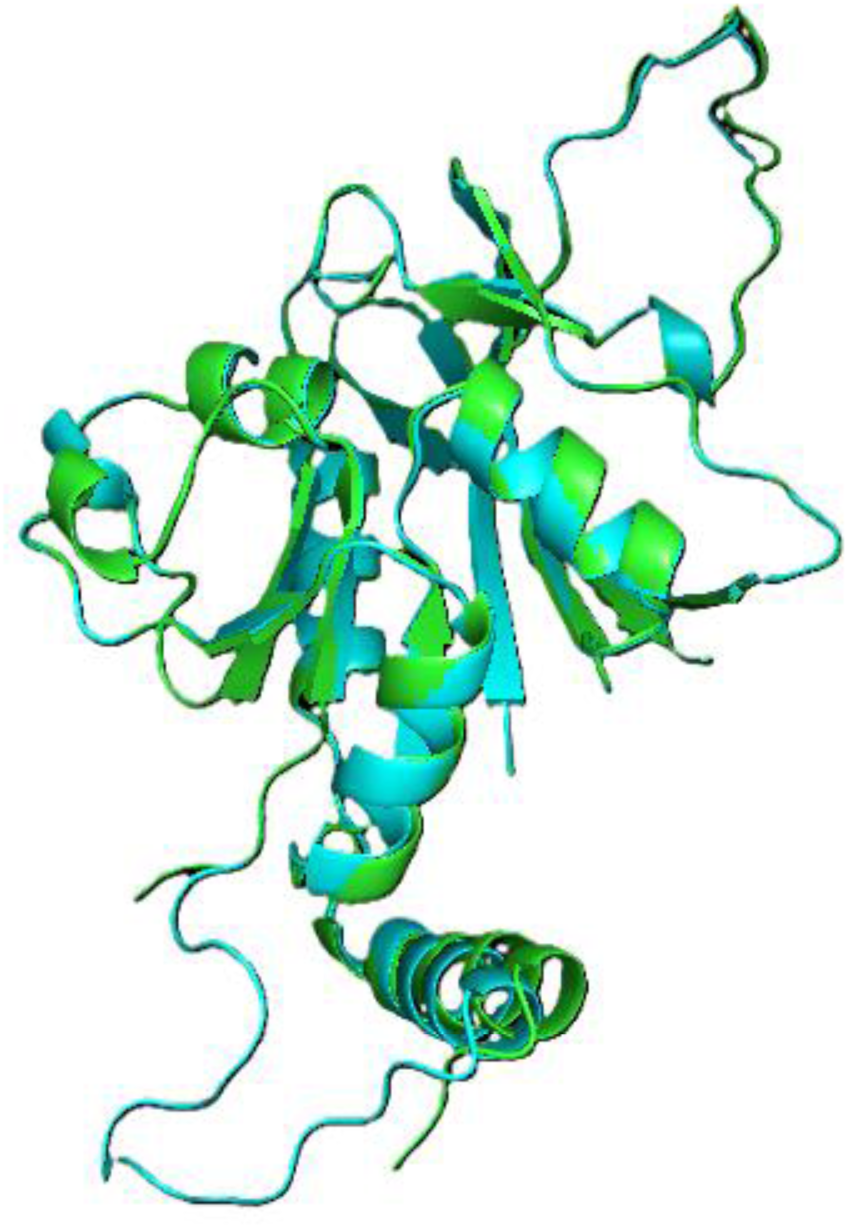
Shows the structural alignment of the modeled DHFR domain (cyan) with the template structure (green) using PyMOL. The RMSD was calculated to assess the structural similarity between the model and the template. The RMSD value of 0.306 Å indicates a good alignment, validating the accuracy of the homology model. This alignment helps confirm that the modeled structure retains the key features and overall topology of the template.

